# Multiplexed BOLD oscillations reveal the interplay of normalization and attention

**DOI:** 10.1101/2025.07.01.662576

**Authors:** Reebal W. Rafeh, Geoffrey N. Ngo, Lyle E. Muller, Ali R. Khan, Ravi S. Menon, Marieke Mur, Taylor W. Schmitz

## Abstract

Monkey electrophysiology has linked attention to divisive normalization, yet noninvasive evidence in humans remains limited. We use frequency-tagged fMRI to isolate visual cortical populations that simultaneously encode multiple competing inputs. We show that responses of these sites are suppressed during inattention and enhanced during attention – consistent with the normalization model, which predicts that attention selectively disinhibits competing inputs – offering a noninvasive translational bridge to study fine-grained computations underlying attentional selection.

## Main

Our visual environment contains a large amount of competing information that exceeds the processing capacity of the brain ^1^. Normalization provides a predictive framework for how the brain responds under such conditions: when multiple stimuli are presented simultaneously, the neural responses to each are suppressed due to inhibitory interactions among competing representations ^1,2^. This competition is resolved by top-down attentional mechanisms that enhance the processing of behaviorally relevant inputs ^1–3^.

However, translating the normalization framework to the human brain has been challenging due to limitations in non-invasive neuroimaging methods ^4^. In the present study, we leveraged frequency-tagging in combination with functional magnetic resonance imaging (ft-fMRI) to examine how attention biases spatial competition between stimuli, and whether its influence conforms to the predictions of normalization. The frequency-tagging approach allows the simultaneous tracking of distinct feature-tuned neural populations within the same cortical area by isolating their synchronized responses to different stimulation frequencies. This enables the identification of individual cortical vertices that respond selectively to a single input (singular vertices) versus those that respond to multiple inputs (multiplexing vertices). The presence of heterogeneous frequency synchronization within multiplexing vertices indicates functionally distinct neural subpopulations at the sub-vertex scale—providing a novel readout of the fine-grained cortical organization of feature tuning.

A critical prediction of the normalization model is that the suppressive drive on feature-selective neurons arises from the pooled activity of nearby populations with overlapping tuning profiles ^2,3^. In the context of ft-fMRI, multiplexing vertices—those showing synchronization to multiple input frequencies—reflect the combined activity of co-activated, feature-tuned subpopulations. As a result, these vertices should exhibit greater suppressive interactions and thus lower baseline response amplitudes under conditions of stimulus competition. Crucially, normalization also predicts that attention selectively mitigates this suppression by disinhibiting subpopulations tuned to the attended stimulus. This disinhibition should yield a more pronounced increase in signal at multiplexing vertices relative to singular vertices when attention is engaged.

To test these hypotheses, we used a ft-fMRI design to delineate visual cortical populations that are feature-tuned to the spatial locations of two spatially competing stimuli. To do so, we first localized vertices exhibiting robust in-phase synchronization at the frequencies (F1 or F2) of either of the presented frequency-tagged checkerboard wedge stimuli ^5^ (Fig. 1a; Supplementary Fig. 1a). Next, we separated vertices exhibiting multiplexing responses from those exhibiting singular responses according to the thresholding criteria described in ^5,6^ (Fig.1a, Supplementary Fig. 1). Because our stimulus was designed a priori to drive BOLD oscillations in location-tuned ventral visual areas, we focus our subsequent analyses on regions of interest (ROIs) anatomically constrained to areas V1v-hV4 ^7^. Consistent with prior work showing that population receptive field sizes increase moving up the visual hierarchy ^8^, we found that the proportion of multiplexing vertices increased from V1v to hV4 (Kruskal-Wallis test: H(2)_F1_ = 7.33, p_F1_= 0.026; H(2)_F2_ = 3.35, p_F2_ = 0.188) (Fig. 1b, Supplementary Table 1).

**Figure 1:**
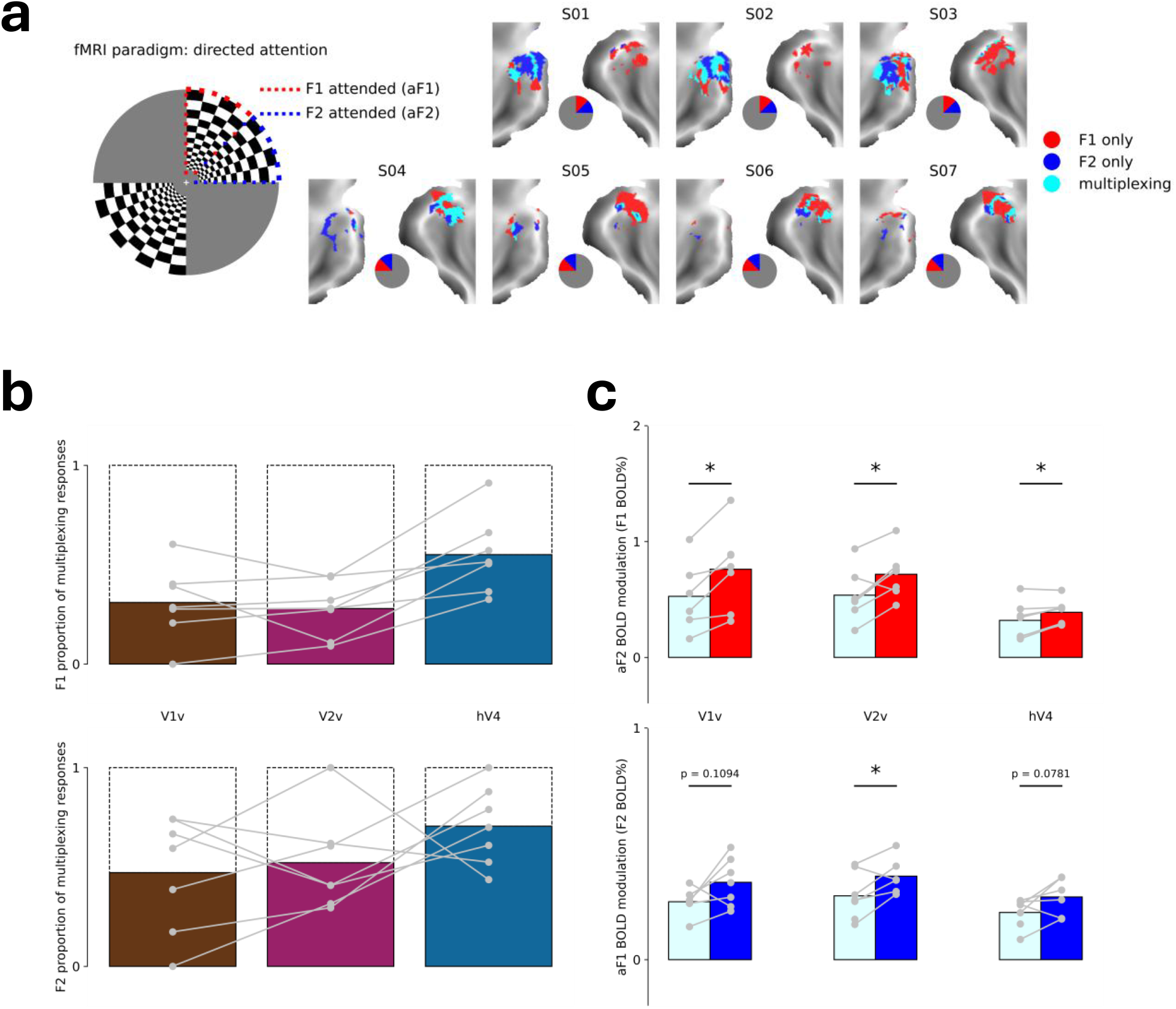
**(a)** Stimulus configuration and vertices localized on the ventral visual surface exhibiting multiplexing (cyan) or singular responses to each of the stimulation frequencies F1 (red) and F2 (blue). Central insets indicate the spatial location at which each of the frequency-tagged checkerboard wedge stimuli were presented for each participant. **(b)** Proportion of multiplexing vertices relative to the total number of localized vertices in each ROI at F1 (top) and F2 (bottom) **(c)** Oscillatory BOLD amplitude during the condition in which the frequency of interest was unattended (aF2:top; aF1; bottom) in vertices exhibiting multiplexing and singular responses. Statistical significance was determined using a Wilcoxon signed-rank test (*P_one-tailed_ < 0.05).

We next examined the amplitude of BOLD oscillations in multiplexing and singular vertices at each of our frequencies of interest during the condition in which the stimulus frequency was unattended (i.e. F1_BOLD%_ during attend-F2; F2 _BOLD%_ during attend-F1) (Fig. 1c). Consistent with normalization, multiplexing vertices exhibited significantly lower amplitudes of oscillatory BOLD responses to each of the competing input frequencies than singular vertices, indicating stronger suppression (Wilcoxon signed-rank test: p_F1 one-tailed_ = 0.016, D_F1_ = 1.17; p_F2 one-tailed_ = 0.016, D_F2_ = 1.52). This effect was replicable across experimental sessions (Supplementary Fig. 2a), and was present across ROIs along the ventral visual surface (Fig. 1c; Supplementary Fig. 2b) (Wilcoxon signed-rank test V1: p_F1 one-tailed_ = 0.016, D_F1_ = 0.710; p_F2 one-tailed_ = 0.11, D_F2_ = 0.972; V2: p_F1 one-tailed_ = 0.016, D_F1_ = 0.841; p_F2 one-tailed_ = 0.047, D_F2_ = 0.939; hV4: p_F1 one-tailed_ = 0.023, D_F1_ = 0.509; p_F2 one-tailed_ = 0.078, D_F2_ = 0.927).

Having established that multiplexing vertices exhibit suppressed responses to each of the competing input frequencies relative to singular vertices, we next asked how directed attention affects the oscillatory signal amplitude in each of these populations. To address this, we computed changes in the amplitude of BOLD oscillations in multiplexing and singular populations when the stimulus was attended relative to when it was unattended (ΔF1_BOLD%_: attend-F1 - attend-F2; ΔF2_BOLD%_: attend-F2 - attend-F1).

We first examined the effect of attentional modulation on the oscillatory BOLD amplitudes of multiplexing and singular vertices separately for F1 (aF1 - aF2; Fig. 2a) and F2 (aF2 - aF1; Fig. 2b). For both frequencies, we found that attention significantly enhanced the amplitude of the multiplexing (Wilcoxon signed-rank test: p_F1 one-tailed_ = 0.0156, D_F1_ = 1.499; p_F2 one-tailed_ = 0.008, D_F2_ = 1.479), but not the singular vertices (Wilcoxon signed-rank test: p_F1 one-tailed_ = 0.711, D_F1_ = -0.261; p_F2 one-tailed_ = 0.531, D_F2_ = 0.040). We also directly compared the magnitude of attentional modulation between the multiplexing and singular vertices for F1 [(aF1_multiplexing_ -aF2_multiplexing_) - (aF1_singular_ -aF2_singular_)] and F2 [(aF2_multiplexing_ -aF1_multiplexing_) - (aF2_singular_ -aF1_singular_)]. For both frequencies F1 and F2, and across experimental sessions (Supplementary Fig. 3a), we observed that the amplitude of BOLD oscillations was significantly more enhanced in multiplexing relative to singular vertices (Wilcoxon signed-rank test: p_F1 one-tailed_ = 0.008, D_F1_ = 1.951; p_F2 one-tailed_ = 0.008, D_F2_ = 1.546).

**Figure 2:**
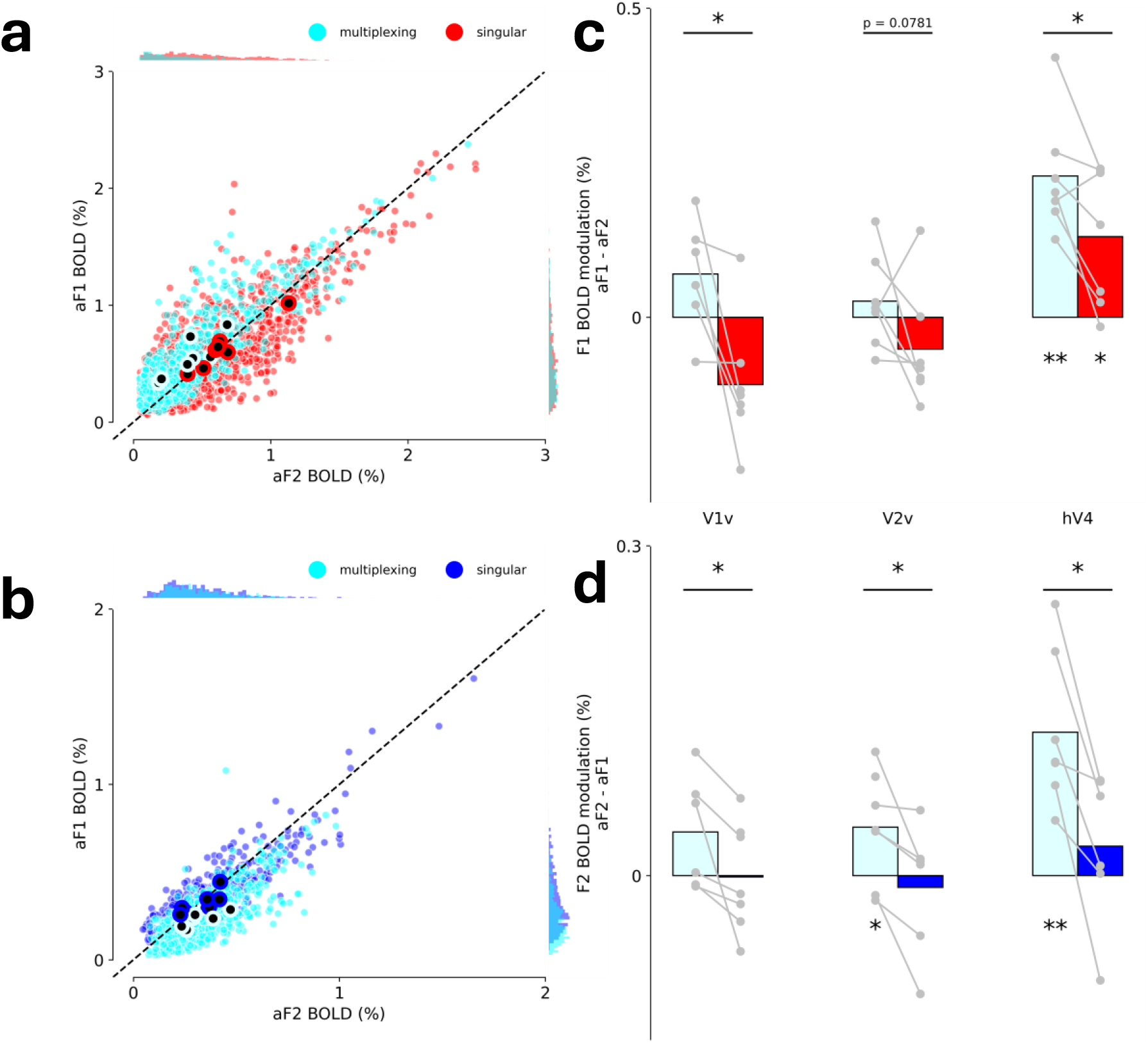
**(a)** The relationship of aF1 (y-axis) to aF2 (x-axis) across all vertices of all participants for the multiplexing (cyan) and singular (red) vertices (small closed circles), and the vertex-wise averages for each participant (closed large circles) at F1. Dots along the diagonal indicate no difference in BOLD modulation between aF1 and aF2. (b) The relationship of aF2 (x-axis) to aF1 (y-axis) across all vertices of all participants for the multiplexing (cyan) and singular (blue) vertices at F2. Representation of the data follows the same convention as in (a). **(c)** Attention-driven modulation of the amplitudes of multiplexing and singular oscillatory BOLD responses for F1 (aF1-aF2). Gray dots represent the average attentional modulation across multiplexing or singular vertices within an ROI for each participant. **(d)** Attention-driven modulation of the amplitudes of multiplexing and singular oscillatory BOLD responses for F2 (aF2-aF1). Representation of the data follows the same convention as in (c). Statistical significance was determined using a Wilcoxon signed-rank test (\**P*_one-tailed_ < 0.05, \*\**P*_one-tailed_ < 0.01).

We next examined whether the effects of attention on oscillatory BOLD responses change as a function of processing stage moving up the visual cortical hierarchy (Fig. 2c,d; Supplementary Fig. 4). Overall, a clear pattern emerges where directed attention increasingly enhances the amplitude of BOLD oscillations in multiplexing and singular vertices moving up the visual hierarchy, with the strongest effects observed in hV4 (Multiplexing: Kruskal-Wallis test H(2)_F1_ = 11.4, p_F1_= 0.003; H(2)_F2_ = 6.64, p_F2_ = 0.036; Singular H(2)_F1_ = 10.9, p_F1_= 0.004; H(2)_F2_ = 1.31, p_F2_ = 0.520; Supplementary Table 2). However, the effects of attention were more pronounced in multiplexing as compared to singular vertices at all stages of the visual processing hierarchy at both frequencies F1 (Fig. 2c) and F2 (Fig. 2d; Supplementary Fig. 3b) (Wilcoxon signed-rank test V1: p_F1 one- tailed_ = 0.016, D_F1_ = 1.83; p_F2 one-tailed_ =0.016, D_F2_ = 0.802; V2: p_F1 one-tailed_ = 0.078, D_F1_ = 0.898; p_F2 one-tailed_ = 0.016, D_F2_ = 0.985; hV4: p_F1 one-tailed_ = 0.023, D_F1_ = 0.944; p_F2 one-tailed_ = 0.016, D_F2_ = 1.476). These results reproduce the known relationship between attentional modulation and receptive field size ^9,10^, and explicitly link this relationship to the strength of competitive interactions among feature-tuned populations at sub-vertex resolution (multiplexing>singular).

Our results show that multiplexing vertices exhibit reduced oscillatory BOLD amplitudes compared to singular vertices during inattention (Fig. 1), and that attention disproportionately enhances BOLD amplitudes in these multiplexing populations (Fig. 2), consistent with predictions from the normalization model. According to this model, suppression arises from mutual inhibition between feature-tuned subpopulations, such that multiplexing vertices—which respond to multiple features simultaneously—exhibit stronger suppressive interactions. Attention is hypothesized to resolve this competition by selectively enhancing the responses of subpopulations tuned to the attended input, thereby disinhibiting them and shifting the push-pull balance in their favor (Fig. 3a).

**Figure 3:**
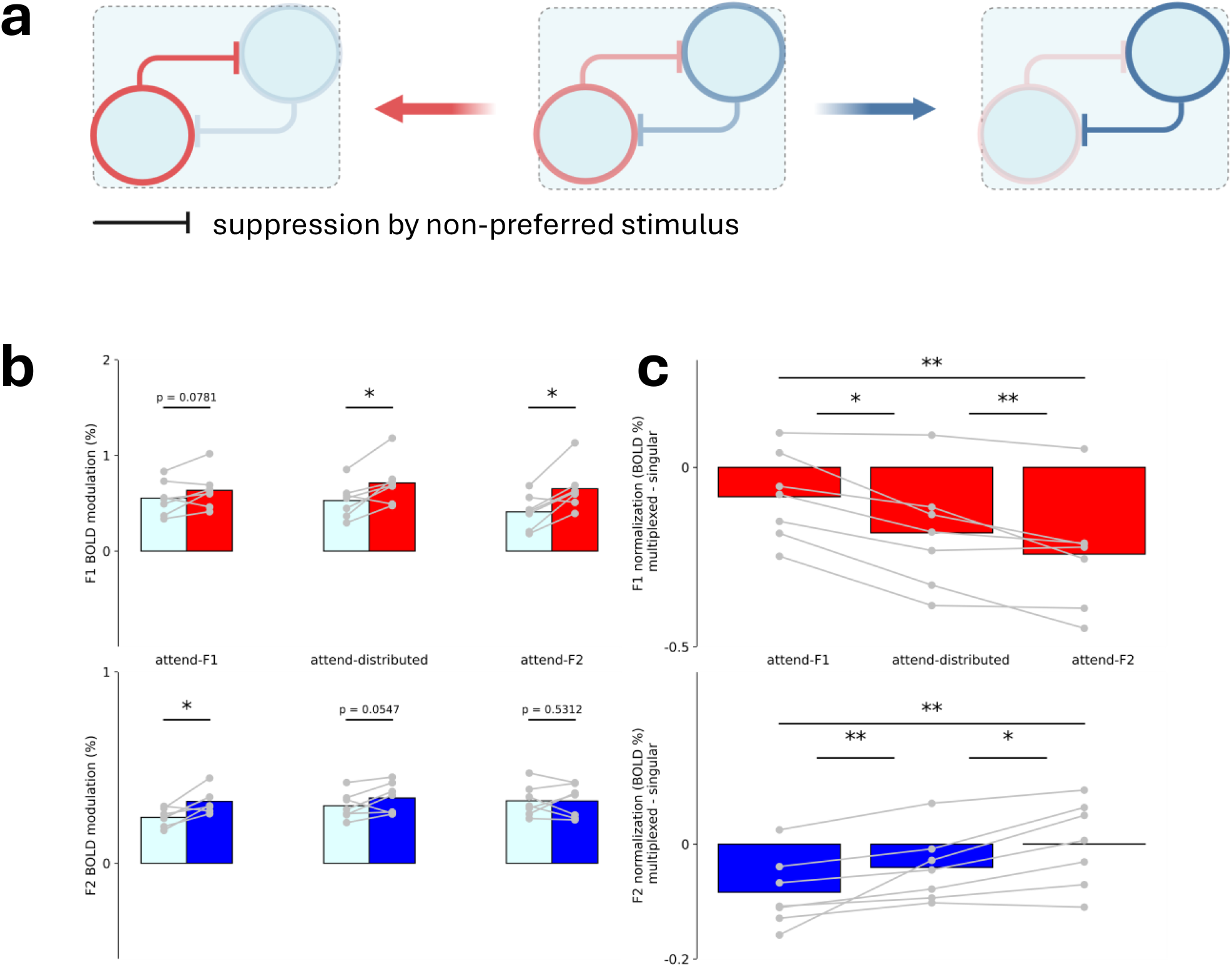
**(a)** Normalization framework schematic describing the push-pull dynamic between populations differentially tuned to F1 or F2 within multiplexing vertices as a function of attention **(b)** Differences in the amplitude of the BOLD signal between vertices exhibiting singular versus multiplexing responses with increasing levels of attention. **(c)** Changes in the strength of normalization with increasing levels of attention. Attention reduces the difference in amplitude between the BOLD oscillations exhibited in multiplexing vertices and those exhibiting singular responses at both F1 (attend-F1 vs. attend-distributed: p_F1 one-tailed_ = 0.008, D_F1_ = 0.716; attend-distributed vs. attend-F2: p_F1 one-tailed_ = 0.023, D_F1_ = 0.375; attend-F1 vs. attend-F2: p_F1 one-tailed_ = 0.008, D_F1_ = 1.122; Wilcoxon signed-rank test versus zero: attend-F2: p_F2 one-tailed_ = 0.016, D_F2_ = -1.511; attend-distributed: p_F2 one-tailed_ = 0.016, D_F2_ = -1.168; attend-F1: p_F2 one-tailed_ = 0.078, D_F2_ = -0.668) and F2 (attend-F2 vs. attend-distributed: p_F2 one-tailed_ = 0.016, D_F2_ = 0.375; attend-distributed vs. attend-F1: p_F2 one-tailed_ = 0.008, D_F2_ = 0.716; attend-F2 vs. attend-F1: p_F2 one-tailed_ = 0.008, D_F2_ = 1.122; Wilcoxon signed-rank test versus zero: attend-F2: p_F2 one-tailed_ = 0.531, D_F2_ = 0.007; attend-distributed: p_F2 one-tailed_ = 0.054, D_F2_ = -0.675; attend-F1: p_F2 one-tailed_ = 0.016, D_F2_ = -1.356) Statistical significance was determined using a Wilcoxon signed-rank test (\**P*_one-tailed_ < 0.05, \*\**P*_one-tailed_ < 0.01).

To test this mechanism, we examined how attentional modulation varied across conditions of increasing attentional bias: attend-F1, distributed attention, and attend-F2. As predicted, attention progressively reduced the amplitude difference between multiplexing and singular vertices, consistent with reduced suppression within the multiplexing population (Fig. 3b). This pattern was observed for both stimulation frequencies and supported by significant one-tailed Wilcoxon signed-rank tests comparing attentional conditions (Fig. 3c caption).

These results support the hypothesis that attention operates by modulating the strength of normalization-based suppression within multiplexing vertices, i.e., fine-grained spatially interdigitated populations tuned to the driving stimuli, preferentially disinhibiting the population best tuned to the attended stimulus. This disinhibition enhances the amplitude of synchronized BOLD responses, effectively biasing the outcome of local population competition in favor of behaviorally relevant inputs.

Our findings demonstrate that attention modulates neural competition in the human visual cortex through mechanisms consistent with divisive normalization ^1,2,11^. By combining frequency-tagging with fMRI, we isolated the oscillatory BOLD responses of distinct feature-tuned populations organized at a fine-grained spatial scale exceeding vertex resolution. Vertices exhibiting multiplexing responses to both stimuli showed attenuated amplitudes during inattention—consistent with normalization-driven suppression—and exhibited greater enhancement under attention, consistent with selective disinhibition. These results provide evidence that attention acts locally to bias the outcome of competitive interactions among neural populations tuned to different visual features.

Traditional fMRI studies of normalization have provided important insights by comparing BOLD responses across different stimulus configurations or attentional states ^10,12^. While powerful, these methods infer competitive interactions indirectly and may conflate normalization-related effects with other sources of variability tied to stimulus context or task demands. In contrast, ft-fMRI enables the simultaneous measurement of responses to multiple competing inputs within the same voxel, allowing us to resolve feature-specific neural dynamics at finer spatial and temporal scales. By synchronizing neural populations to distinct stimulation frequencies, we can track the effects of attention on each input independently, revealing how suppression and disinhibition unfold within multiplexing cortical sites in real time.

This approach brings human neuroimaging into closer alignment with invasive electrophysiological studies of normalization in non-human primates, where neuronal responses to simultaneously presented, feature-tuned stimuli can be dissociated in single-unit recordings. ft-fMRI provides an analogous population-level readout, but at a whole-brain scale and with noninvasive accessibility—offering a powerful translational tool for bridging animal models and human cognition. By resolving competition at the level of co-localized subpopulations, this method opens new opportunities for testing computational theories of attention and for evaluating how normalization dynamics may be altered in clinical populations characterized by attentional dysfunction or excitatory-inhibitory imbalance.

## Methods

### Participants

Eight right-handed participants (4 female) with normal or corrected-to-normal vision participated as paid volunteers after giving informed written consent. Their ages ranged from 20-31 years old (M = 26.25, SD = 3.49). Participants were compensated at a rate of CAD$15 per hour for their participation in the study. Participants also received an additional CAD$0.05 for each correct response during the behavioral task. One female participant was excluded from further analyses due to large gaze position deviations from fixation throughout the experiment (gaze area across experimental runs > 5 degrees^2^). All experimental procedures were approved by the Health Sciences Research Ethics Board at Western University.

### Stimuli & Task

We generated the experimental stimuli and tasks using PsychoPy ^13^. Stimuli were projected at 60 Hz onto a screen (resolution of 1024×768 pixels) fixed at the back of the scanner bore from a frame rate synchronized stimulus laptop. Participants viewed the projected visual stimuli via a mirror fitted into the head coil placed in front of the eyes (total distance to screen is 84 cm; 10° visual angle).

We presented participants with a pair of wedge stimuli occupying one visual field quadrant and another pair of radial wedge stimuli occupying equal spatial portions of the opposing visual field quadrant. Each wedge was part of a checkerboard pattern. The wedge stimuli extended from 1.5° to 10° eccentricity. Participants initiated each experimental run via a button press. During a 14s baseline period at the beginning of each experimental run, we used a red line(s) to cue our participants to covertly attend to the wedge(s) in one visual field quadrant. Following cue offset, participants had to covertly attend to and detect transient color changes (lasting for 500 ms) at either (directed attention task) or both (distributed attention task) oscillating wedge stimuli presented in either the upper right or upper left visual field quadrant. Oscillating wedge stimuli were sine-squared modulated at 0.125 Hz and 0.2 Hz, and exhibited a 12 Hz counterphase flicker.

Color changes only occurred when the oscillating stimuli were at above 33% luminance contrast. Color change inter-stimulus-interval was adjusted for the frequency of the oscillating wedges. For the stimulus oscillating at 0.125 Hz, color changes occurred at 0.5-2 s pseudo-random intervals drawn from a uniform distribution. For the stimulus oscillating at 0.2 Hz, color changes occurred at 0.5-1.25 s pseudo-random intervals drawn from a uniform distribution. Color changes during the directed attention conditions had a 50% chance of occurring on each oscillation of the wedge. The participant had 750 ms to respond to the color change occurrence via button press with the right index finger. If the response did not occur within the response interval, the response was recorded as a false alarm. If the participant did not respond at all before the next color change appeared, this was recorded as a missed trial. Color change intensity was continuously adjusted on each experimental run using an adaptive staircase procedure (QUEST) ^14^ to achieve a response accuracy of approximately 80%, where response accuracy was calculated as the number of correct responses divided by the total sum of correct responses, false alarms and missed trials. QUEST parameters: ß (slope of psychometric function) = 3.5; δ (probability of incorrect response under ideal conditions) = 0.01; γ (probability of false alarm) = 0.05; Reference stimulus color change = 40% of maximum contrast.

Each participant performed a total of sixteen 219s runs of each of the three experimental conditions; ‘distributed-attention’, ‘attend-F1’, ‘attend-F2’, over the course of four experimental sessions (4 runs of each condition per session). We counterbalanced the location of the stimuli in either the upper right or upper left visual hemifield across participants (4 participants per location).

### MRI acquisition

MRI data were acquired on a Siemens Magnetom 7T machine (MAGNETOM MRI Plus, Siemens Healthineers, Erlangen, Germany) at the Center for Functional and Metabolic Mapping using a Siemens 32-channel whole-head array for the anatomical T1-weighted images and a custom occipital-parietal 32-channel head coil for the functional T2/T2*-weighted images.

A T1-weighted anatomical MP2RAGE scan was acquired during the first study visit for registration purposes and to enable cortical surface projection (TR 6000 ms, TE 2.74 ms, TI1 800 ms, TI2 2700 ms, FOV 240 mm, flip angle 1 4°, flip angle 2 5°, 0.7 mm isotropic voxels, 224 slices, AP phase encoding direction, in-plane acceleration factor 3, slice partial Fourier 6/8). fMRI scans were collected during each of the four study visits using a multiband T2*-weighted echo-planar imaging sequence covering an oblique slab centered on the ventral occipital lobe (TR 250 ms, TE 20 ms, FOV 208 mm, flip angle 30°, 2.5 mm isotropic voxels, 16 slices, RL phase encoding direction, no in-plane acceleration, and multi-band acceleration factor 4) with 880 volumes acquired per scan for a total acquisition time of 220s. T2-weighted anatomical images (TR 2,000 ms, TE 20 ms, FOV 256 mm, flip angle 30°, 2 mm isotropic voxels, 80 slices, RL phase encoding direction, in-plane acceleration factor 2, and multi-band acceleration factor 4) were acquired in the middle of each study session to facilitate the registration between session-specific slab fMRI data and the T1-weighted anatomical image.

Data were converted from dicom to nifti brain imaging data structure (BIDS) format using heudiconv (v0.11.3) ^15^.

### MRI data processing

MRI data was preprocessed using a pipeline developed with *Nipype*, which used the following software packages: ANTS, Freesurfer, FSL, AFNI, workbench, and customized Nipype workflows adapted from smriprep and fmriprep ^6^. Existing Nipype workflows from fmriprep were customized to integrate session-specific whole-brain EPIs and run-specific EPIs into the registration workflow.

To preprocess each fMRI dataset, slice-timing correction, followed by single-shot interpolation to T1w space, surface resampling to 32k vertex fsLR surfaces (fsLR 32k surfaces), and nuisance regression were performed. fMRI data was regressed using 24 motion parameters (six HMC parameters and their 1st and 2nd order first derivatives), high pass filter regressors (<.01 Hz), and the mean WM and CSF signal to minimize the effects of scanner drift, motion and other non-neural physiological noises. Each fMRI run was truncated between 59 and 219 seconds (or 45 seconds after the oscillatory stimulus begins and when the stimulus ends) to account for transient hemodynamic effects associated with the onset of continuous oscillatory stimulation, and to ensure that the run duration provided the frequency resolution necessary for precise phase estimation at the frequency of both oscillating wedge stimuli. Since our stimuli had periods of 8s (0.125 Hz) and 5s (0.2 Hz), we were able to obtain 20 and 32 full cycles, respectively, at our frequencies of interest from the resulting pre-processed and noise-corrected 160s time course.

Details pertaining to resampling of individual fMRI runs to fsLR 32k surfaces are as follows: The participant-specific T1-weighted (T1w) structural image was skull-stripped using SynthStrip and then processed with smriprep to segment the grey matter (WM), white matter (GM), and cerebrospinal fluid (CSF). Subsequently, the T1w image was registered to the single-band reference (SBRef) image from each scan-specific whole-brain EPI using a boundary-based registration (BBR) cost function. The cortical ribbon mask was transformed into the whole-brain EPI space for each session and utilized to perform a BBR-based registration between the SBRef images of the scan-specific whole-brain EPIs and their corresponding scan-and-run-specific slab EPIs. N4 bias field corrections were performed to each image prior to performing the registrations. Next, after applying low-pass filtering to each fMRI dataset, head motion correction (HMC) was performed using FSL’s MCFLIRT, registering all volumes to their associated run-specific SBRef image. We low-pass filtered the HMC parameters to remove unwanted fluctuations (0.2-0.4 Hz) in the estimated HMC time courses. Note that low-pass filtered data was used solely to generate fluctuation-free HMC transformations, which were then applied to the raw, unfiltered fMRI data. To enable one-shot interpolation of each fMRI run to T1w space, the transformations were concatenated in the following order: HMC to run-specific slab EPI, slab EPI to session-specific whole brain EPI, and whole brain EPI to participant-specific T1w image. Next, the volumetric BOLD data (in T1w space) was resampled onto the participant’s native surface, and subsequently resampled onto a standard fsLR 32k surface.

### Activation thresholding

Preprocessed fMRI runs were grouped for each participant for the localizer condition (distributed-attention task) to generate frequency-encoded brain maps. A Monte Carlo random subsampling approach was used to ensure that the localized frequency-encoded populations were robust against random subsamples. This involved performing 500 iterations of model fitting. In each iteration, preprocessed fMRI runs from a single participant were randomly split into train and test groups (50/50 split). An average fMRI run was computed for each data split, followed by time series normalization to BOLD % change relative to the mean of the time series, and subsequent fitting with a set of frequencies.

To generate a frequency-encoded map, a general linear model was fitted on each iteration using a Fourier basis set at each frequency (F1 = 0.125 Hz; F2 = 0.2 Hz; see *Fitting frequency-encoded populations*). Maps of each frequency were binarized by thresholding with an unadjusted p-value < 0.05 (F-test). These activation maps were aggregated across all 500 iterations by computing the mean, producing a fractional overlap map that indicates the stability of frequency-encoded brain maps across random subsampled iterations. A value of zero means the frequency was not encoded in any of the 500 random subsamples, while a value of one means it was encoded in all 500 subsamples. Based on our previous work ^5,6^, we thresholded frequency-encoded maps at a fractional overlap value of 0.8 (i.e. vertices exhibiting significant frequency-specific modulation in at least 400/500 subsamples).

### Fitting frequency-encoded populations

To model a frequency of interest in the fMRI data, vertex-wise BOLD time series were fitted using a general linear model. The model included sine and cosine regressors associated with each frequency of interest, and an F-test was performed on the combined fit to the sine and cosine regressors at each frequency of interest to test for a frequency-specific response. The amplitude and phase of the frequency-specific response was then determined using the coefficient weights of the sine and cosine components at each frequency. Description of the general linear model is formally described below.

For any given BOLD time series, the general linear model can be expressed as follows:

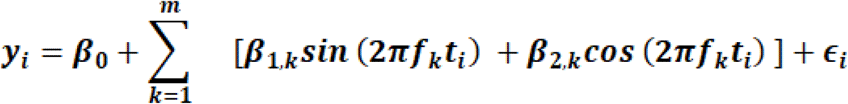

where:

- β_0_ is the intercept term
- β_*1*,*k*_ and β_*2*,*k*_ are the coefficients associated to the sine and cosine terms of the k^th^ frequency *f*_*k*_, respectively
- *f*_*k*_ is the k^th^ frequency of interest
- ϵ _*i*_ is the error term for observation *i*
- *m* is the number of frequencies of interest

The amplitude (*BOLD%*_*k*_) at a frequency of interest was calculated as the root sum squared of the fitted coefficients associated with the Fourier basis of that frequency. This value was then multiplied by 2 to include the negative portion of the oscillatory response in the amplitude (peak-to-trough) estimate. For each participant, task condition, and *k*^*th*^ frequency of interest, we obtained 500 estimates of the response amplitude via the iterative resampling procedure described in *Activation thresholding*. The reported amplitude estimate for each vertex corresponds to the mean amplitude across resamplings of the data.

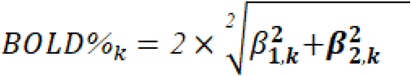

Amplitude,

The phase-offset (ϑ_*k*_) of a frequency of interest was calculated as the arctangent of the Fourier basis coefficients of that frequency. For each participant, task condition, and *k*^*th*^ frequency of interest, we obtained 500 estimates of the phase offset via the iterative resampling procedure described in *Activation thresholding*. The reported phase estimate for each vertex corresponds to the circular mean of the phase estimates across resamplings of the data.

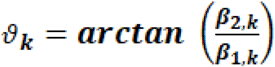

Phase offset,

### Classification of in-phase and anti-phase synchronized responses

To separate vertex-wise data into populations that synchronize in-phase and anti-phase with the stimulus, we fit a von Mises mixture model to the phase offset data estimated from the localizer (distributed-attention task) using the *clustering* algorithm from the *pycircstat2* library. This resulted in two unique phase distributions for each frequency of interest. To determine which of these distributions described in-phase or anti-phase responses, we first convolved the stimulus time course at each frequency with a gamma hemodynamic response function (HRF), and truncated the convolved time course to the period of the analysed data (that is, 59s - 219s). Next, we calculated the phase of the resulting time course to obtain an estimate of the in-phase response predicted by the canonical HRF. We computed the circular distance between each distribution and the predicted HRF phase by obtaining the average circular distance across vertex phase estimates within a distribution and the predicted HRF phase. We then defined vertices falling within the phase distribution with the smallest circular distance to the predicted HRF phase as responding in-phase with the stimulus. Vertices falling within the opposing distribution were defined as responding out of phase (anti-phase) with the stimulus.

### Definition of vertices exhibiting multiplexing and singular responses

To delineate vertices that were feature-tuned to the location of each of our stimuli, we selected vertices exhibiting significant (fractional overlap > 0.8) in-phase synchronization with the stimulus frequencies. Multiplexing vertices were defined as those exhibiting significant in-phase responses at the frequencies of both presented stimuli (F1 and F2), whereas singular vertices were defined as those exhibiting significant in-phase responses at only one of the two frequencies (F1 only or F2 only).

### ROI definition

For each participant, we selected ROIs from a visual surface-based cortical parcellation ^7^. We specifically selected ROIs that were targeted by our stimulation protocol (i.e. those located on the ventral visual surface: V1v, V2v, V3v, hV4). We only included cortical hemisphere-specific data from an ROI in our analysis if we could on average localize at least 5 suprathreshold vertices (fractional overlap > 0.8) across resamplings of the data in that hemisphere. ROI data from visual region V3v was excluded from analyses since we could only localize multiplexing vertices in 3/7 participants in this region.

### Attentional modulation analysis

To obtain an estimate of attentional modulation at our frequencies of interest, we first calculated the difference in BOLD amplitude between the condition in which the frequency of interest was attended relative to when the frequency was unattended across 500 resamplings of the data in each of these conditions (F1 attentional modulation = aF1_BOLD%_ -aF2_BOLD%_; F2 attentional modulation = aF2_BOLD%_ -aF1_BOLD%_). We then averaged the attentional modulation values across estimates to obtain the reported measure of attentional BOLD amplitude modulation for multiplexing (Supplementary Fig. 5) and singular vertices (Supplementary Fig. 6).

### Eye-tracking data acquisition & analysis

Eye tracking was performed using an Eyelink 1000 system during the experiment. Eye tracking data was sampled at 0.5 kHz. Before each experimental session, eye position was calibrated using a 9-point calibration. For each experimental run, we obtained a time series of horizontal and vertical eye positions. Blinks were automatically detected by the Eyelink system and manually selected based on outlier values, and data within ±0.1 seconds of each blink were excised. We detrended recorded eye-positions to account for instrumental drift over the course of the experimental session. We plotted a 2D histogram of horizontal and vertical eye-positions for every experimental run. Eye positions were summarized by fitting a 2D Gaussian probability distribution to the data and calculating the area of the contour that contained 95% of the fitted distribution, and gaze area for each condition was computed as the mean area of the fitted distribution across runs (Supplementary Fig. 7a,b).

### Behavioral data acquisition & analysis

During each experimental run, we recorded participant button press responses to a target detection task (see Stimuli & Task). Participant behavioral performance on the task was determined as the number of correct responses divided by the total sum of correct responses, false alarms and missed trials (Supplementary Fig. 7c).

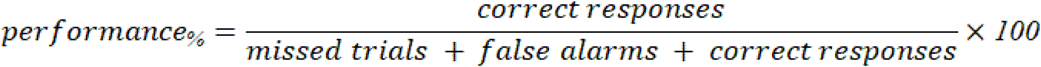

### Statistical analysis

Statistical analysis was performed using the scipy and statsmodels libraries in python. For vertex-wise inference, the statsmodels library was used to fit the GLM to the time series data and perform the F-test to determine the significance of frequency-specific responses. For participant-wise statistical inference, no assumptions were made about the normality of compared data distributions. The scipy library was used to perform the reported non-parametric statistical tests. An alpha level of p < 0.05 was used to assess significance.

## Data & Code Availability Statement

Example data and analysis code will be made available upon the publication of this manuscript, or via reasonable request to the corresponding author.

## References

1. Desimone, R. & Duncan, J. Neural mechanisms of selective visual attention. Annu. Rev. Neurosci. 18, 193–222 (1995).

2. Reynolds, J. H. & Heeger, D. J. The normalization model of attention. Neuron 61, 168–185 (2009).

3. Verhoef, B.-E. & Maunsell, J. H. R. Attention-related changes in correlated neuronal activity arise from normalization mechanisms. Nat. Neurosci. 20, 969–977 (2017).

4. Sejnowski, T. J., Churchland, P. S. & Movshon, J. A. Putting big data to good use in neuroscience. Nat. Neurosci. 17, 1440–1441 (2014).

5. Rafeh, R. W. et al. Attentional enhancement and suppression of stimulus-synchronized BOLD oscillations. bioRxiv (2025) doi:10.1101/2025.01.10.632431.

6. Ngo, G. N. et al. Frequency-tagged fMRI: A platform for fine-grained spatiotemporal analysis of cortical function. bioRxiv (2024) doi:10.1101/2024.12.19.629518.

7. Wang, L., Mruczek, R. E. B., Arcaro, M. J. & Kastner, S. Probabilistic maps of visual topography in human cortex. Cereb. Cortex 25, 3911–3931 (2015).

8. Kay, K. N., Weiner, K. S. & Grill-Spector, K. Attention reduces spatial uncertainty in human ventral temporal cortex. Curr. Biol. 25, 595–600 (2015).

9. Moran, J. & Desimone, R. Selective attention gates visual processing in the extrastriate cortex. Science 229, 782–784 (1985).

10. Kastner, S., De Weerd, P., Desimone, R. & Ungerleider, L. G. Mechanisms of directed attention in the human extrastriate cortex as revealed by functional MRI. Science 282, 108–111 (1998).

11. Maunsell, J. H. R. Neuronal Mechanisms of Visual Attention. Annu Rev Vis Sci 1, 373–391 (2015).

12. Bloem, I. M. & Ling, S. Normalization governs attentional modulation within human visual cortex. Nat. Commun. 10, 5660 (2019).

13. Peirce, J. et al. PsychoPy2: Experiments in behavior made easy. Behav. Res. Methods 51, 195–203 (2019).

14. Watson, A. B. & Pelli, D. G. QUEST: a Bayesian adaptive psychometric method. Percept. Psychophys. 33, 113–120 (1983).

15. Halchenko, Y. O. et al. HeuDiConv -flexible DICOM conversion into structured directory layouts. J. Open Source Softw. 9, 5839 (2024).

