## Supplementary Materials for "Multiplexed BOLD oscillations reveal the interplay of normalization and attention"

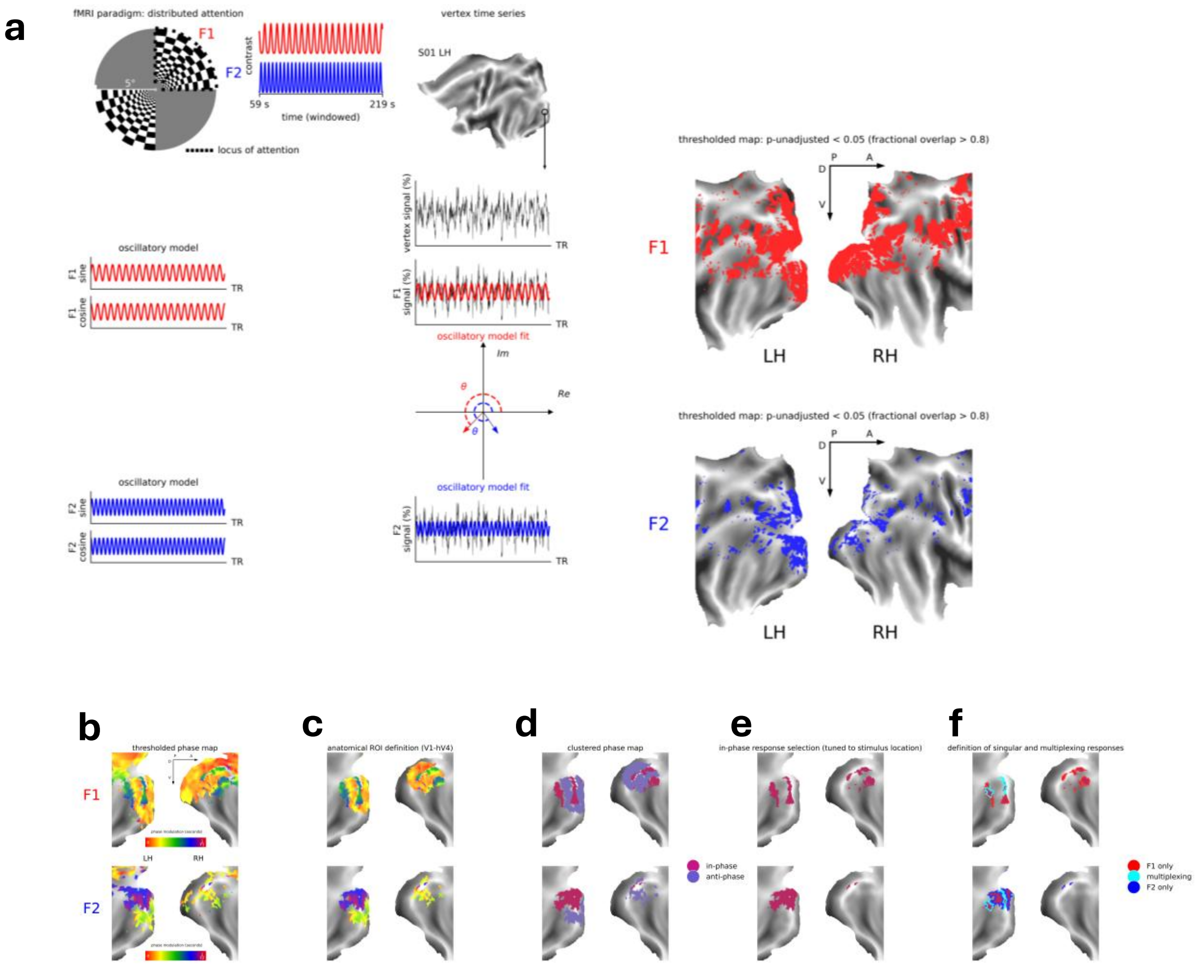


***Supplementary Figure 1: Flowchart schematic showing the localization and definition of feature-tuned vertices exhibiting multiplexing or singular oscillatory BOLD responses***

***(a)*** *Stimulus configuration and BOLD response phase modelling. Participants covertly directed their attention to a pair of checkerboard wedges oscillating at different frequencies (F1 = 0.125 Hz; F2 = 0.2 Hz) in an upper visual field quadrant. For each run, we analyzed the resulting vertex timeseries in a 160 s time window (59s – 219s). A sine and cosine function at each stimulation frequency was fit to the timeseries of each vertex using general linear models (GLM). The fitted sine and cosine functions were then projected onto the complex plane, providing an estimate of BOLD signal amplitude and phase. This procedure was repeated 500 times for each vertex using different resamplings of the data. Right: Maps of vertices exhibiting robust stimulus-synchronized responses at F1 (red:top) and F2 (blue:bottom). We thresholded vertices according to the proportion of resamples exhibiting a significant model fit at our frequencies of interest (F-test; p-unadjusted < 0.05). Vertices exhibiting a significant GLM fit across at least 80% of the random samples (i.e., 400/500 resamples) were determined to be task-driven.****(b)*** *The localized visual cortical phase maps at F1 (top) and F2 (bottom) projected to the flattened surface of a representative participant (S01). The reported phase value for each vertex is the circular mean of phase estimates across resamples of the data. The color bar denotes the response phase scaled to the period of the examined frequency.* ***(c)*** *Anatomical ROI selection along the ventral visual surface for further analysis (V1-hV4)* ***(d)*** *Clustering of response phases into in-phase and anti-phase frequency synchronized vertices* ***(e)*** *Selection of in-phase responses (i.e. vertices tuned to the spatial location of the frequency-tagged stimulus) for further analysis* ***(f)*** *Delineating homogeneously tuned singular vertices exhibiting significant responses to either of the two competing frequency-tagged stimuli (F1: red; F2: blue), and heterogeneously tuned multiplexing vertices exhibiting significant responses to both frequency-tagged stimuli (cyan)*

**a**

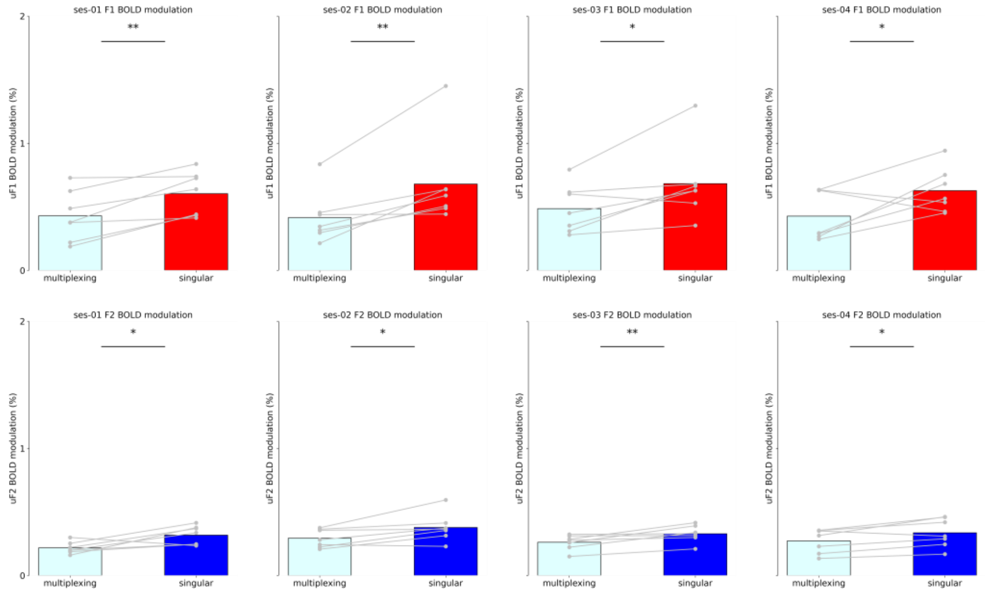


**b**

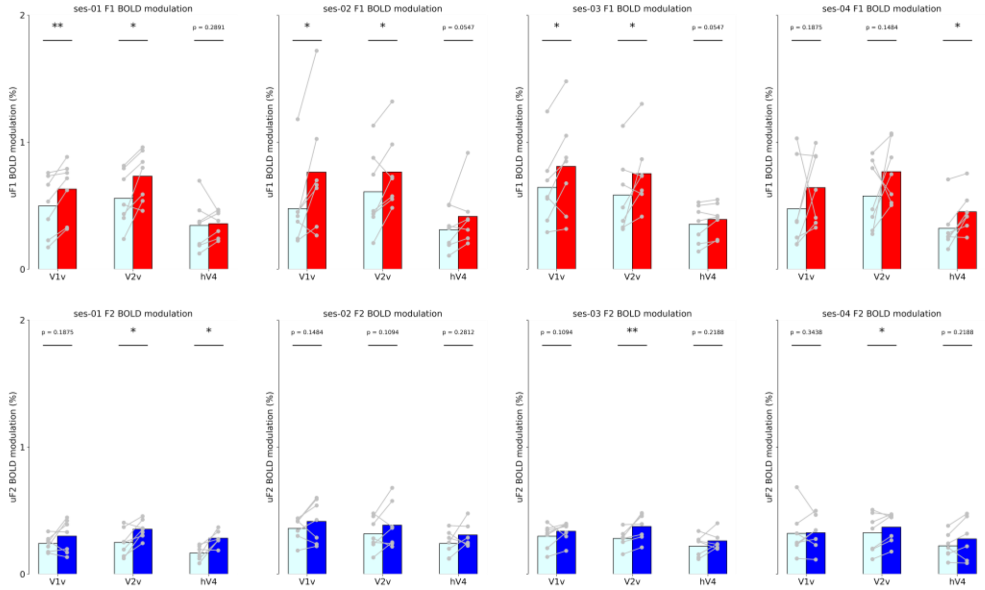


***Supplementary Figure 2: Reliability of normalization effects across experimental sessions***

***(a)*** *Oscillatory BOLD amplitude during the condition in which the frequency of interest was unattended (aF2:top; aF1; bottom) across all localized vertices exhibiting multiplexing (cyan) and singular (F1: red; F2: blue) responses across experimental sessions (columns). Gray dots represent the average BOLD response across multiplexing or singular vertices for each participant.* ***(b)*** *Representation of data follows the same convention as in (a), but for individual visual ROIs. Statistical significance was determined using a Wilcoxon signed-rank test (*$P_{one\text{-}tailed} < 0.05$; **$P_{one\text{-}tailed} < 0.01$)*

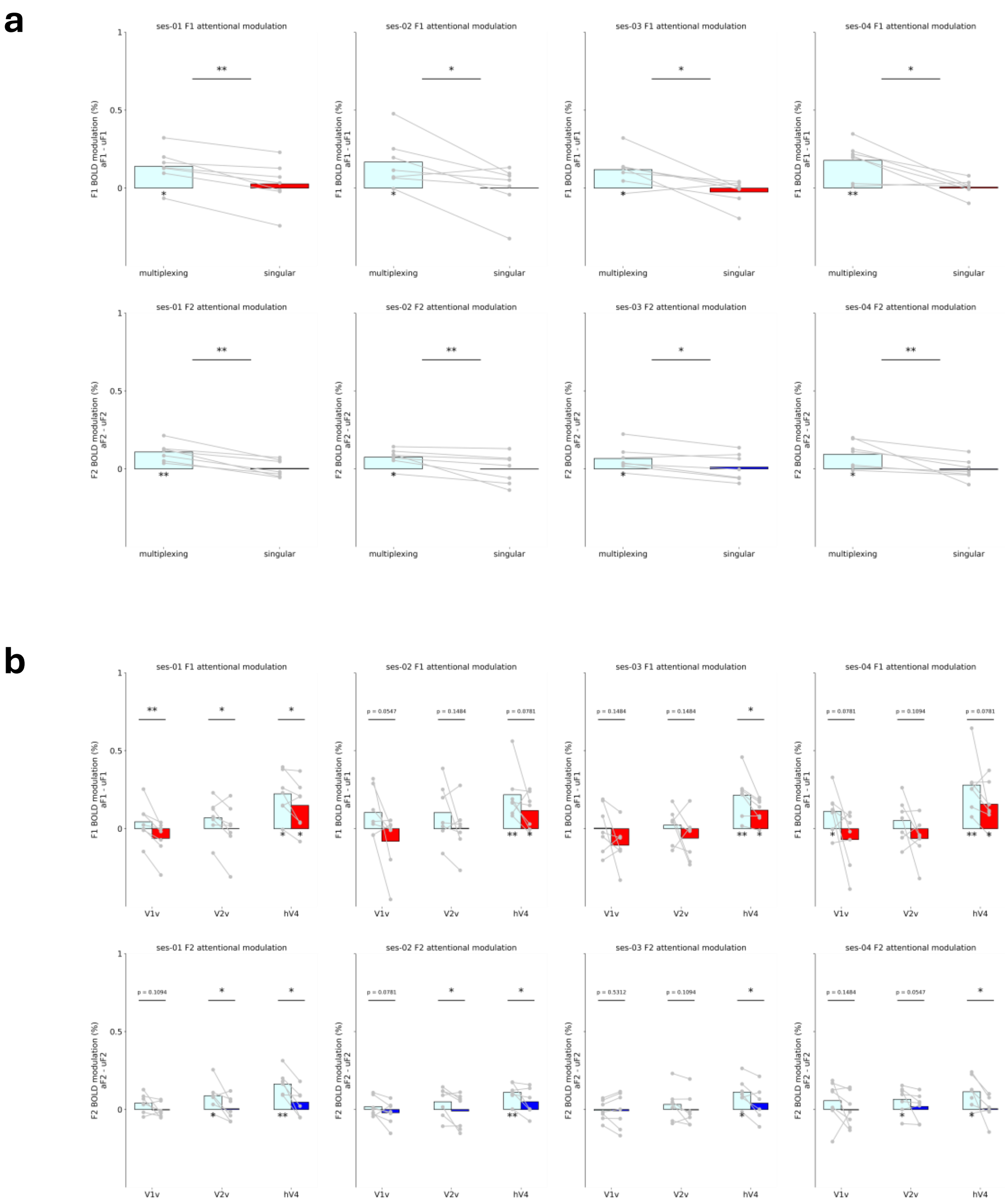


***Supplementary Figure 3: Reliability of attentional modulation effects across experimental sessions***

***(a)*** *Differences in amplitude of BOLD oscillations between multiplexing (cyan) and singular (F1: red; F2: blue) ROIs during the condition in which attention is directed away from the stimulus (top: aF2; bottom: aF1) relative to the condition in which attention is directed towards the oscillating stimulus (top: aF1; bottom: aF2). Gray dots represent the average attentional BOLD modulation value across multiplexing or singular vertices for each participant.* ***(b)*** *Representation of data follows the same convention as in (a), but for individual visual ROIs. Statistical significance was determined using a Wilcoxon signed-rank test (*$P_{one\text{-}tailed} < 0.05$*; **$P_{one\text{-}tailed} < 0.01$*).*

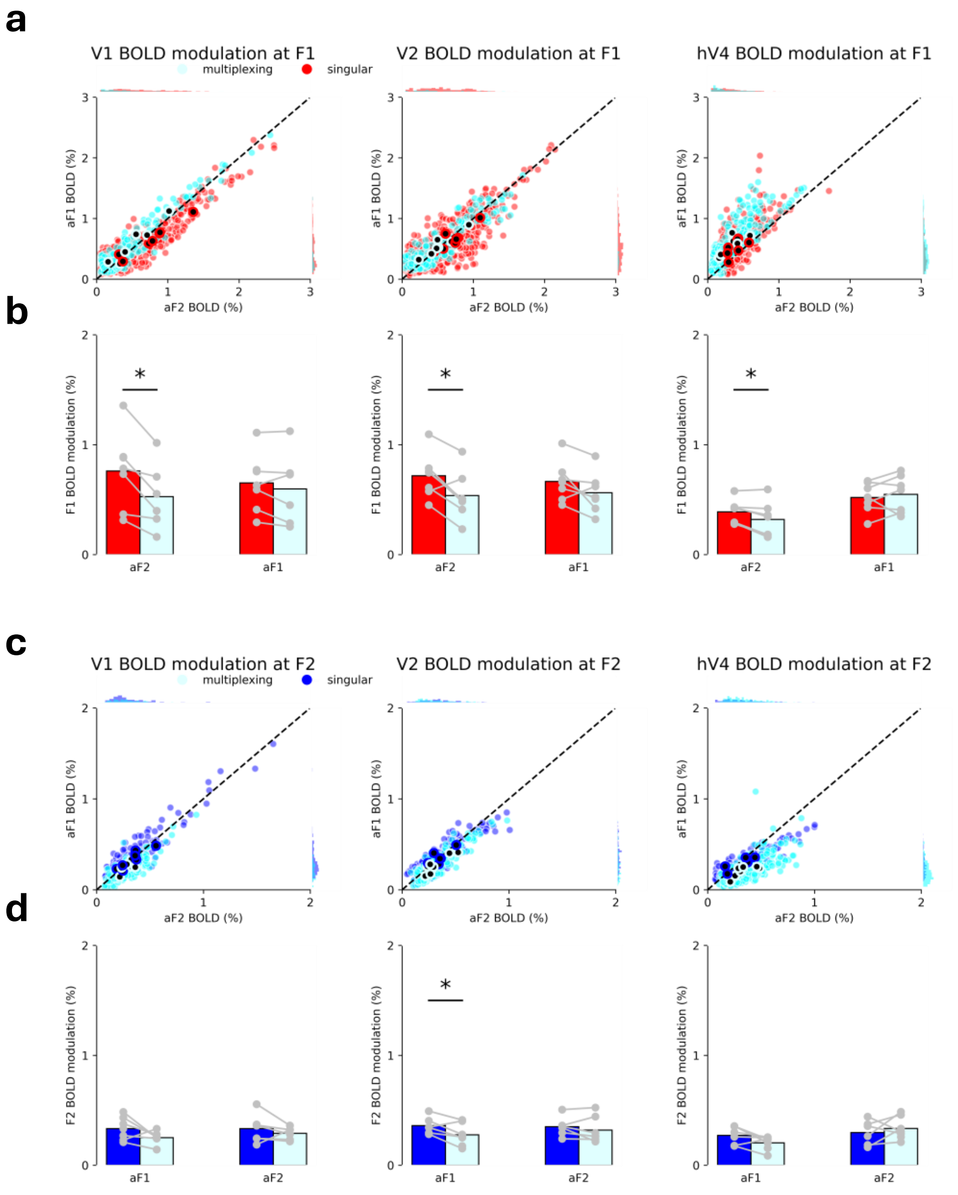


***Supplementary Figure 4: Attentional modulation of oscillatory BOLD amplitude of vertices exhibiting multiplexed and singular responses in different visual ROIs***

*__(a)__ The relationship of aF1 (y-axis) to aF2 (x-axis) for visual ROI (columns) vertices of all participants for the multiplexing (cyan) and singular (red) vertices. Dots along the diagonal indicate no difference in BOLD modulation between aF1 and aF2. __(b)__ Differences in amplitude of BOLD oscillations between multiplexing (cyan) and singular (red) ROIs during the condition in which attention is directed away from the stimulus (aF2) relative to the condition in which attention is directed towards the oscillating stimulus (aF1). __(c)__ The relationship of aF2 (x-axis) to aF1 (y-axis) for visual ROI (columns) vertices of all participants for the multiplexing (cyan) and singular (blue) vertices. Representation of the data follows the same convention as in (a). __(d)__ Representation of the data follows the same convention as in (b). Statistical significance determined using a Wilcoxon signed-rank test (*$P_{one\text{-}tailed} < 0.05$).*

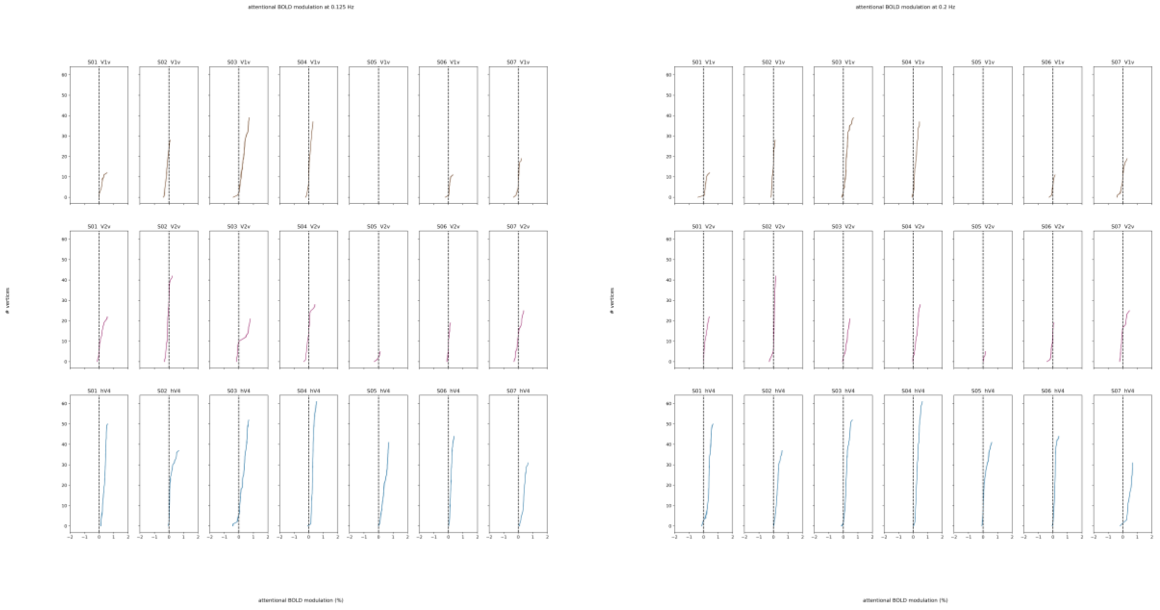


***Supplementary Figure 5: Consistency of attentional modulation of multiplexing vertices across visual ROIs***

*Individual participant data showing the consistency of attentional modulation estimates in multiplexing vertices across participants (columns) and ROIs (rows) at F1 (left) and F2 (right). We examined the estimated attentional modulation for each vertex across resamplings of the data (500 repetitions), and we show the 95% CI across resamplings. The 95% CI indicates that estimates of attentional modulation in multiplexing vertices were consistent across resamplings of the data for each participant and analyzed ROI.*

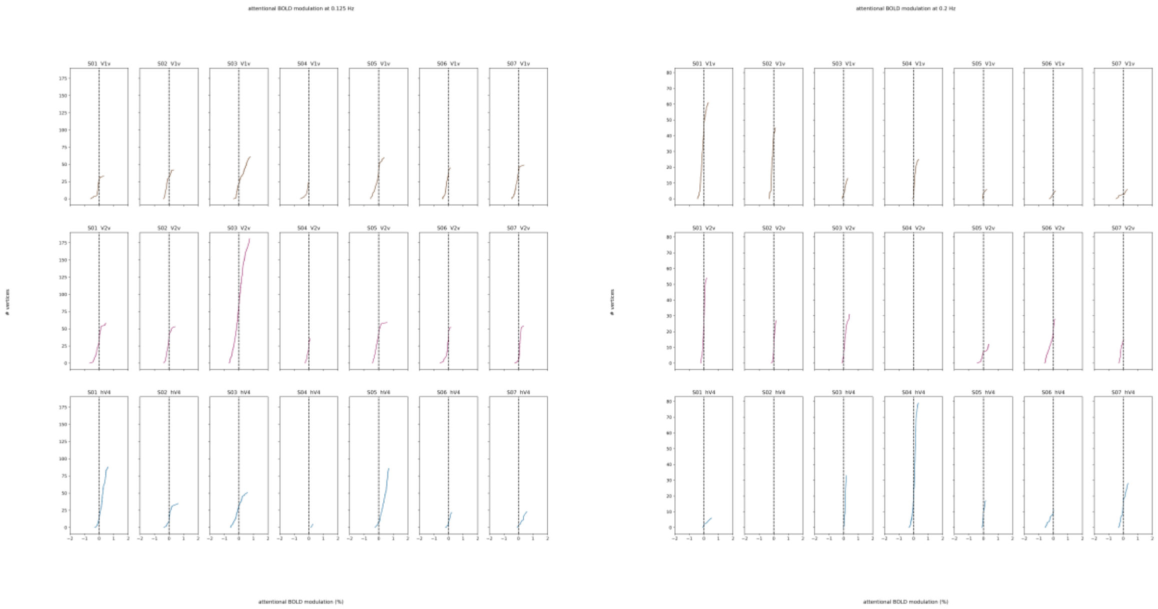


***Supplementary Figure 6: Consistency of attentional modulation of singular vertices across visual ROIs***

*Individual participant data showing the consistency of attentional modulation estimates in singular vertices across participants (columns) and ROIs (rows) at F1 (left) and F2 (right). We examined the estimated attentional modulation for each vertex across resamplings of the data (500 repetitions), and we show the 95% CI across resamplings. The 95% CI indicates that estimates of attentional modulation in singular vertices were consistent across resamplings of the data for each participant and analyzed ROI.*

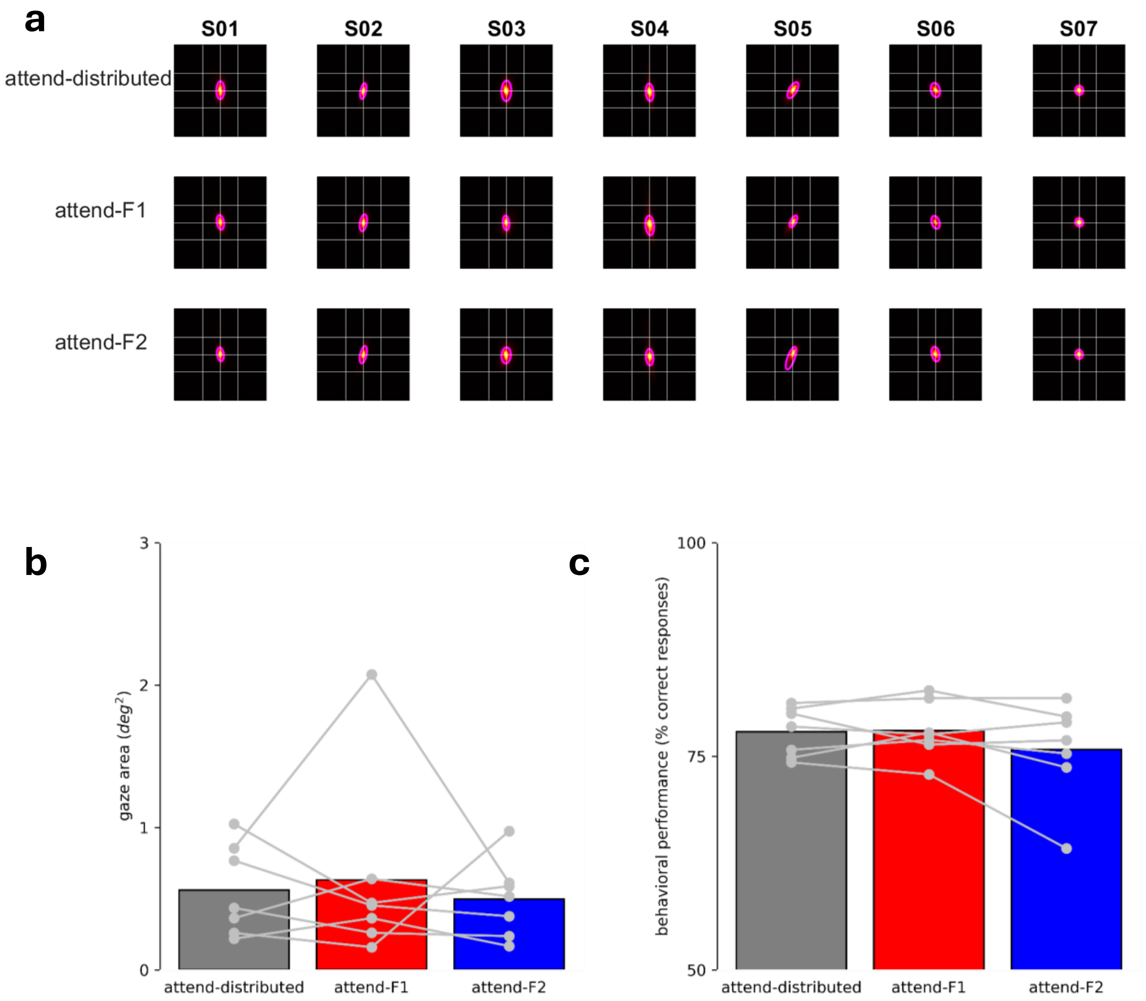


***Supplementary Figure 7: Participant behavioral performance across experimental conditions***

***(a)*** *Eye positions remain consistently near the center of the display. Eye-tracking data is summarized by displaying the heatmaps and fitted gaussian distribution of the aggregated data for each participant (columns) and experimental condition (rows). The grid lines indicate the radius of the central fixation region. The magenta contour lines indicate the area of the gaussian mixture model that contains 95% of the fitted data.* ***(b)*** *The gaze contour areas are comparable between experimental conditions across participants (K-W ANOVA; H(2) = 0.052; $p_{two\text{-}tailed}$=0.974). Data points indicate the participant mean gaze contour area across runs for an experimental condition. Data points are within the area of the central fixation region (~1.8 $deg^2$). Individual participants showed comparable gaze-areas between experimental conditions across runs (one-way ANOVA: $p_{two\text{-}tailed} > 0.05$ for 7/7 participants).* ***(c)*** *Participant behavioral performance was comparable between experimental conditions across participants (K-W ANOVA; H(2) = 0.586; $p_{two\text{-}tailed}$ = 0.746). Data points indicate the participant mean behavioral performance across runs for an experimental condition. These data show that the adaptive staircase procedure maintained behavioral performance at ~80%. Individual participants showed comparable behavioral performance between experimental conditions across runs (one-way ANOVA: $p_{two\text{-}tailed} > 0.05$ for 6/7 participants).*

***Supplemental Table 1: Test statistic = Wilcoxon signed rank tests with statistical null of no difference in the proportion of multiplexing vertices in an ROI across visual ROIs.***

| **Figure** | **ROI Comparison** | **Frequency** | **Test statistic** | **P-value** |
|---|---|---|---|---|
| 1b | V1 vs. V2 | F1 | Rank = 13 | $p_{two\text{-}tailed} = 0.9375$ |
| **1b** | **V1 vs. hV4** | **F1** | **Rank = 0** | $\mathbf{p_{two\text{-}tailed} = 0.0156}$ |
| **1b** | **V2 vs. hV4** | **F1** | **Rank = 0** | $\mathbf{p_{two\text{-}tailed} = 0.0156}$ |
| 1b | V1 vs. V2 | F2 | Rank = 12 | $p_{two\text{-}tailed} = 0.8125$ |
| 1b | V1 vs. hV4 | F2 | Rank = 9 | $p_{two\text{-}tailed} = 0.4688$ |
| 1b | V2 vs. hV4 | F2 | Rank = 7 | $p_{two\text{-}tailed} = 0.2969$ |

***Supplemental Table 2: Test statistic = Wilcoxon signed rank tests with statistical null of no difference between the amplitudes of BOLD oscillations between the attended and unattended frequency, and alternative of greater oscillatory BOLD amplitude during attended relative to unattended condition.***

| Figure | Vertex population | Attended Frequency | Visual ROI | Test statistic | P-value |
|---|---|---|---|---|---|
| 2c | multiplexing | F1 | V1 | Rank = 18 | $p_{one\text{-}tailed}$ = 0.078 |
| 2c | multiplexing | F1 | V2 | Rank = 19 | $p_{one\text{-}tailed}$ = 0.234 |
| **2c** | **multiplexing** | **F1** | **hV4** | **Rank = 28** | **$p_{one\text{-}tailed}$ = 0.008** |
| 2c | singular | F1 | V1 | Rank = 2 | $p_{one\text{-}tailed}$ = 0.984 |
| 2c | singular | F1 | V2 | Rank = 7 | $p_{one\text{-}tailed}$ = 0.891 |
| **2c** | **singular** | **F1** | **hV4** | **Rank = 27** | **$p_{one\text{-}tailed}$ = 0.016** |
| 2d | multiplexing | F2 | V1 | Rank = 16 | $p_{one\text{-}tailed}$ = 0.156 |
| **2d** | **multiplexing** | **F2** | **V2** | **Rank = 25** | **$p_{one\text{-}tailed}$ = 0.039** |
| **2d** | **multiplexing** | **F2** | **hV4** | **Rank = 28** | **$p_{one\text{-}tailed}$ = 0.008** |
| 2d | singular | F2 | V1 | Rank = 14 | $p_{one\text{-}tailed}$ = 0.531 |
| 2d | singular | F2 | V2 | Rank = 11 | $p_{one\text{-}tailed}$ = 0.5 |
| 2d | singular | F2 | hV4 | Rank = 15 | $p_{one\text{-}tailed}$ = 0.219 |
